# Metabuli: sensitive and specific metagenomic classification via joint analysis of amino-acid and DNA

**DOI:** 10.1101/2023.05.31.543018

**Authors:** Jaebeom Kim, Martin Steinegger

## Abstract

Current metagenomic classifiers analyze either DNA or amino-acid (AA) sequences. DNA-based methods have better specificity in distinguishing well-studied clades, but they have limited sensitivity in detecting under-studied clades. AA-based methods suffer the opposite problem. To tackle this trade-off, we developed Metabuli for a joint analysis of DNA and AA using a novel k-mer, *metamer*. In benchmarks, Metabuli was simultaneously as specific as DNA-based methods and as sensitive as AA-based methods. In the CAMI2 plant-associated dataset, Metabuli covers 99% and 98% of classifications of state-of-the-art DNA-based and AA-based classifiers, respectively. Metabuli is available as free and open-source software for Linux and macOS at metabuli.steineggerlab.com.

---

Metagenomics allows studying microbial communities by analyzing DNA or RNA sequences directly taken from various environments. Some studies aim to reveal evolutionary distant organisms (e.g., in the soil (1), ocean (2) and hydrothermal vent sites (3)). Others, in the clinical field, focus on detecting pathogens and emerging strains in samples from patients (4), public spaces (5), and wastewater (6).

Identifying the origin of metagenomic reads is performed by searching for similar regions in reference sequences. One way to detect the similarity is to calculate local alignments between the read and the reference as in MMseqs2 Taxonomy (7) and MEGAN CE (8). Alternatively, alignment-free methods were introduced for faster classification. For instance, k-mer-based tools extract fixed-length k-mers from queries and references and matches them. Another type, FM-index-based tools utilize the Burrows-Wheeler transformation of the references to query (9, 10) k-mer matches of flexible length.

Metagenomic classifier needs two contrasting capabilities: 1) specificity for high-resolution classification of well-studied clades and 2) sensitivity to detect under-studied species based on known relatives in a database.

However, current tools suffer an inherent trade-off problem between specificity and sensitivity depending on the sequence type they utilize: DNA or amino-acids (AAs) (11–13). DNA-based tools have better specificity as they exploit point mutations to differentiate strains. AA-based tools leverage the higher conservation of AA sequences for better sensitivity to detect homology between novel organisms and their relatives in the reference, although it limits resolving close taxa.

As a partial countermeasure, classifiers that are particularly well-suited to the research context need to be selected (11–13). However, metagenomic samples are a mixture of well- and under-studied taxa, the specificity-sensitivity trade-off inevitably restricts full sample characterization.

To address this trade-off problem, we introduce Metabuli, a method that jointly analyzes DNA sequences and their AA translation to achieve both specificity and sensitivity simultaneously (Fig. 1 and Supp. Fig. 1). In benchmarks comprising simulated reads, Critical Assessment of Metagenome Interpretation 2 (CAMI2 (15)) datasets, as well as real-world metagenomes, Metabuli consistently demonstrated top performance while DNA- and AA-based tools had fluctuating performance depending on the distance between the queried organisms and available references in the database.

**Fig. 1.**
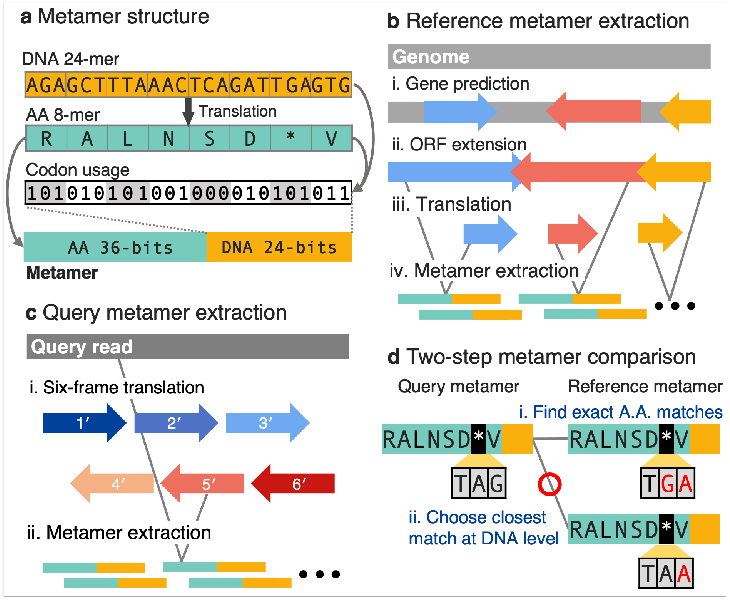
Metabuli’s workflow. **a)** A DNA fragment of 24 nucleotides is translated into eight AAs, which are encoded as an integral value encoded within 36 bits. Each AA has 1-6 synonymous codons, thus requiring three bits to store which one is seen in the fragment. **b)** Metabuli predicts ORFs in a genome using Prodigal and extends them to cover intergenic regions. The extended ORFs are used to extract reference metamers. **c)** Metabuli scans each read in six translational frames to extract query metamers. **d)** The metamers are compared first to find exact AA matches and subsequently to choose the closest one at the DNA level.

To enable the joint analysis of DNA and AA sequences, Metabuli utilizes a novel k-mer structure, *metamer*, encoding a 24 nucleotide-long fragment (eight codons) in 60 bits. Its translation to AAs is encoded by 36 bits, and its codons - by 24 bits. Since an AA is coded by at most six codons, three bits per AA suffice to indicate which one is seen. This joint-encoding is more efficient compared to individual encoding, requiring only 2/3 of the bits.

During database creation, Metabuli predicts open reading frames (ORFs) using Prodigal (16). Each ORFs is extended to cover intergenic regions to cover the whole genome, these regions are often missed in methods utilizing only coding sequences (12, 13). Notably, Metabuli only stores metamers up to one-third of the length of contigs. This is in contrast to AA classifier kAsA (17), which involves storing all k-mers from six frames of the entire genomes, leading to a sixfold increase in size. In addition, Metabuli’s reference metamer list is also shortened by removing metamers that are redundant within each species.

To classify each read, Metabuli computes query metamers from each read and its six-frame translations, which are carried through stop codons. These are compared to reference metamers to find perfect AA matches for sensitivity; among them, matches of the lowest DNA Hamming distance are selected for specificity. Metabuli can quickly calculate the distance with a pre-computed distance matrix designed for metamers. The selected matches are analyzed to score candidate taxa and to classify (Supp. Fig. 2). In this process, Metabuli-P (precision mode) uses score thresholds to reduce false positive and over-confident classifications (Methods, Supp. Fig. 3).

To compare the performance of Metabuli to state-of-the-art classifiers, we conducted inclusion and exclusion tests using prokaryotes and viruses (Fig. 2a-d). In inclusion tests, we evaluated specificity, i.e., how well a classifier can distinguish between reads from closely related organisms at lower taxonomic ranks. Thus, query (sub)species were present in the reference as well as their siblings. In contrast, exclusion tests evaluated sensitivity, i.e., the ability to classify reads from a novel (sub)species based on sequences of its siblings, so the query (sub)species was removed from the reference.

**Fig. 2.**
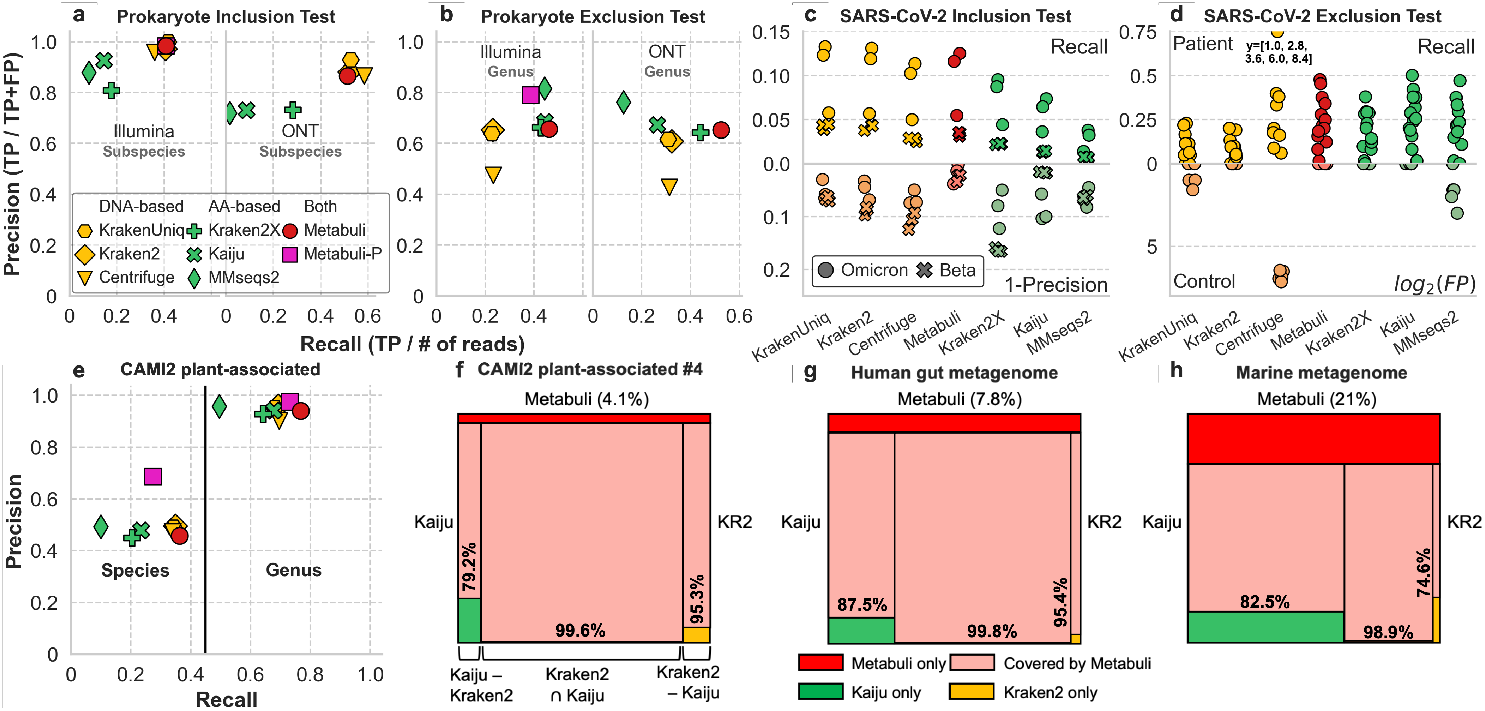
Benchmark results. **a-b) GTDB benchmarks** GTDB genomes and taxonomy were used. Simulated short (Illumina) and long (ONT) reads were used. a) Reads were simulated from genomes present in databases. b) Not the queried species but their sibling species were contained in databases. **c-d) Pathogen detection tests**. RNA-seq reads from COVID-19 patients were classified. c) The reference included five SARS-CoV-2 variants and Viral RefSeq (14), and the reads from patients infected with either omicron or beta variant were queried. Classifications to correct and incorrect variants were counted as TP and FP, respectively. d) Viral RefSeq, excluding SARS-CoV-2, served as the reference. RNA-seq reads from controls (bottom) and patients (top) were classified. Classifications to Sarbecovirus, the LCA of SARS-CoV-1 and 2, were counted as TP for patient samples and as FP for controls. Centrifuge classified more reads as SARS-CoV-2 than the estimated total number of SARS-CoV-2 reads. Recall values for such cases were denoted. **e-f) CAMI2 plant-associated dataset**. e) GTDB genomes and the CAMI2-provided taxonomy were used for database construction. Classifications for the CAMI2-provided plant-associated reads were evaluated. f) The relationship among TP (at genus level) sets of Kaiju, Kraken2, and Metabuli for one of the plant-associated samples. **g-h) Real metagenomes** Human gut (g) and marine (h) metagenomic reads were classified using databases of (a). The proportion of Metabuli-only area in the union of the three tools is denoted in parentheses. f-h) The area is proportional to the number of reads within each panel.

Depending on the purpose of each test, we measured the precision (P) and recall (R) at different ranks. In inclusion tests, we measured them at the (sub)species rank, and in exclusion tests - at the rank of the lowest common ancestor (LCA) of each query and its siblings. When measuring at a certain rank, unclassified reads as well as reads classified at higher ranks were considered false negatives (FNs) to penalize less informative classifications. Meanwhile, classifications at lower ranks climbed up the taxonomy to the rank of measurement. Afterward, classifications to the correct or to the wrong taxon were counted as true positives (TPs) and false positives (FPs), respectively.

First, we designed a short read benchmark using the Genome Taxonomy Database (GTDB) (15). In the inclusion test, where reads were simulated from 1,191 species that had at least two subspecies in the database (19% of all species), DNA-based methods classified more reads to correct subspecies than AA-based ones (Fig. 2a). AA-based methods classified less than 18% of the reads, about half of what DNA-based methods could, also with lower precision. However, in the exclusion test, where reads were simulated from 367 species that were removed from the database, AA-based tools performed better(Fig. 2b). They classified about twice as many reads as DNA-based tools into the correct genus with better precision (R > 0.4 for AA-based, R < 0.25 for DNA-based). These results clearly demonstrate the pros and cons of DNA- or AA-based tools.

Next, we conducted similar tests using simulated long reads. Again, DNA-based tools outperformed AA-based ones in the inclusion test. In contrast, only one AA-based tool, Kraken2X, exceeded DNA-based ones in the exclusion test. Kraken2X ignores frame information, while the other AA-tools are sensitive to frame-shifting indel errors that are more frequent in long reads (18).

Remarkably, only Metabuli achieved top-level performance in all the inclusion and exclusion tests using short and long reads. In the inclusion test, Metabuli performed as well as all DNA-based methods and outperformed all AA-based tools (Fig. 2a). Its performance was more similar to that of DNA-based tools in species rank (Extended Data Fig. 1). Moreover, in the exclusion tests (Fig. 2b), Metabuli achieved the best recall with competent precision with both short and long reads. Since Metabuli scores candidate taxa using matches from multiple frames like Kraken2X, it could be robust to the frequent indels of long reads. Metabuli-P was tested only with short reads for which it is optimized, and it was the second most precise tool with comparable R to AA-based tools in the short read exclusion test.

Next, the classifiers were evaluated using real SARS-CoV-2 data for two main pathogen detection tasks: strain identification and emerging pathogen discovery, both were performed in inclusion and exclusion tests (Fig 2c-d). In the inclusion test, RNA-seq reads from six COVID-19 patients were examined to identify the culprit variant when its genome was present in databases. In contrast, only SARS-CoV-1, but not 2, was provided in the reference databases of the exclusion test.

DNA-based tools classified more reads to the culprits than the AA-based tools in the inclusion test. In the exclusion test, however, the best-performing DNA-based tool, KrakenUniq missed two patient samples and made FP hits in three controls. On the other hand, AA-based tools outperformed DNA-based tools in the exclusion test, detecting up to twice as much SARS-CoV-2. However, they were worse at deciphering the specific culprit strain in the inclusion test.

Here as well, it was only Metabuli and Metabuli-P that showed robust performance in both tests. Their performance was similar, so only Metabuli is depicted in Fig. 2c-d. In the inclusion test, Metabuli classified a comparable number of reads to the culprits as DNA-based methods, even outperforming Centrifuge (10). Moreover, it achieved the best precision, classifying fewer reads to incorrect variants. In the exclusion test, it detected as many SARS-CoV-2 reads as the AA-based Kaiju (9) without any FP hits in the controls.

Next, we sought to challenge the classifiers to identify reads from datasets that contained organisms varying in their query-to-database distances, as would be the case in many real-world studies. To that end, we used query datasets from CAMI2: strain-madness, marine, and plant-associated, which have different query-to-database distances.

On the strain-madness data (Extended Data Fig. 2a), Metabuli and DNA-based tools performed better than AA-based tools. In the marine benchmark (Extended Data Fig. 2b), which contains reads with larger query-to-database distances, the gap in recall became smaller and all tools showed similar precision (*>*0.93). For the plant-associated data with the largest query-to-database distance, tools of both types showed similar performance while Metabuli had the best sensitivity. To investigate this result, we analyzed Metabuli with respect to the genus-level TP sets of the best-performing AA- and DNA-based tools, Kaiju and Kraken2. We found that Metabuli covered 99.6% of their intersection, 79.2% of Kaiju−Kraken2, and 95.3% of Kraken2−Kaiju, which implies that Metabuli successfully joins DNA- and AA-based classifications. Moreover, about 4.1% of the total reads were correctly classified only by Metabuli. Across the three CAMI2 datasets, Metabuli-P progressively improved in precision with the growing diversity of data, with the largest improvement on the plant-associated data set (Extended Data Fig. 2).

Next, we compared Kraken2, Kaiju, and Metabuli using real metagenomic data from well-studied (human gut) and under-studied (marine) environments. As real reads have no ground-truth labels, we compared the proportion of reads classified by each tool. For the human gut data (Fig. 2g), Kraken2 and Kajiu respectively classified 50% and 65% of the total. However, their classified proportion dropped significantly to 30% and 12% as query-to-database distance increased in the marine data set (Fig. 2h). On both data sets, Metabuli could classify the largest number of reads, covering 83-88% of Kaiju−Kraken2, 75-95% of Kraken2−Kaiju, and *>*98.9% of Kaiju ∩ Kraken2.

Finally, we compared the speed, RAM usage, and database size in the prokaryote benchmarks (Supp. Table 1). All tools took less than ten minutes except for MMseqs2 Taxonomy, which spent *>*100 minutes. Of all, Kraken2X was the fastest and used the least RAM, also having the smallest database. Notably, because Metabuli is designed to utilize a user-specified size of RAM, it can classify reads against any size database as long as it fits in the machine’s hard disk. We demonstrated this feature by measuring performance under various configurations. Metabuli was even able to complete the tasks on a notebook with just 8 GiB RAM and 8 threads (Extended Data Table 1). Even though Metabuli stores both DNA and amino acid sequences, its database size was about 1.5 times that of Kraken2’s probabilistic database.

In summary, Metabuli achieves high specificity and high sensitivity simultaneously by utilizing metamers to jointly analyze sequences at both DNA and AA levels. In benchmarks, only Metabuli showed robust state-of-the-art performance, while other tools sacrificed either sensitivity or specificity depending on their type and the benchmark scenario. The results demonstrate the transformative potential of Metabuli for diverse research contexts. Metabuli allows specific classifications for reads from well-studied species while not losing sensitivity for under-studied organisms. At last, Metabuli is open-source software, and ready-to-use binaries and pre-computed databases are provided (Supp. Tables).

## Data availability

Archives and accession numbers of public data used in the benchmarks are listed in the Supplementary Table. The short and long reads used in prokaryote inclusion and exclusion tests are available at https://doi.org/10.5281/zenodo.8084522.

## Code availability

Metabuli is GPLv3-licensed open-source software. The source code and binaries for Metabuli can be downloaded at metabuli.steineggerlab.com. The scripts used for benchmarks are available at https://github.com/jaebeom-kim/metabuli-analysis.

## Acknowledgements

The authors wish to thank Eli Levy Karin from ELKMO for the scientific feedback and careful reading of the manuscript; Johannes Söding for discussions on metamer encoding; Milot Mirdita for the usability improvement of the software; Sebastian Jaenicke for voluntary examination of the software; and Minjae Kim for feedback on the manuscript.

M.S. acknowledges support from the National Research Foundation of Korea (grants 2019R1A6A1A10073437, 2020M3A9G7103933, 2021R1C1C102065, and 2021M3A9I4021220), the Samsung DS Research Fund, and the Creative-Pioneering Researchers Program through Seoul National University.

## Author contributions

J.K. and M.S. designed the research, developed the software, performed analysis, and wrote the manuscript.

## Competing interests

The authors declare no competing interests.

## Methods

### Simulated read generation

To simulate paired-end short reads used in synthetic benchmarks, we used the *mason*_*simulator* module of Mason2 (20). The reads were 150 nt in length and included simulated errors at rates of 0.11% for mismatches, 0.005% for insertions, and 0.005% for deletions. These error rates were based on the performance of the NovaSeq 6000 sequencer. As with Mason2’s default settings, the mismatch probability at the beginning and end of the reads was set to 0.5% and 0.22%. When provided to MMseqs2 Taxonomy, simulated reads were concatenated with ‘NN’ as it does not support paired-end reads. In the case of long reads, we used PBSIM3 (21) to simulate reads of Oxford Nanopore Technologies with options; `--strategy wgs --method errhmm --errhmm ERRHMM-ONT.model --depth 3`.

### GTDB

The GTDB was used for several benchmarks as well as for the calibration of Metabuli-P as it provides phylogenetically consistent taxonomy based on genomic distance measures. For these, we started with a subset of GTDB R202 consisting of 258,406 genomes from 47,894 species clusters. We used the *GTDB*_*metadata*_*f ilter*.*R* module in the pipeline Struo (22) to obtain a list of 22,973 genomes that were assembled at the level of complete genome or chromosome, had CheckM completeness *>*90, and had CheckM contamination *<*5. The filtered genomes were downloaded using Struo’s *genome*_*download*.*R* module, and 22,819 successfully downloaded genomes of 6,186 species were used. NCBI-style taxonomy dump files for the GTDB were generated by *gtdb*_*to*_*taxdump* (23) module. The proteome corresponding to each genome was computed by Prodigal with default settings.

### Metabuli: Database creation

Metabuli builds a reference database of computed metamers from nucleotide sequences following the procedure below (Supp. Fig. 1a-e).

#### ORF prediction and extension

Metabuli utilizes Prodigal for ORF prediction in reference sequences. To enhance the prediction process’s efficiency, we implemented three optimizations. 1) Metabuli bins reference sequences by species in separate FASTA files, then it trains Prodigal once for each species using the longest sequence of the species’ bin before predicting genes. This approach significantly reduces training time, considering the presence of multiple assemblies for a single species. 2) We narrowed down the calculation range of Prodigal’s dynamic programming during both the training and prediction steps. While this adjustment may cause Prodigal to miss very long genes, it effectively reduced runtime by half in tests performed on an *Escherichia coli* genome. 3) We parallelized the training and prediction processes by distributing jobs for species bins across multiple threads, further accelerating computation. After the gene prediction, genes that are fully nested in longer ones are removed. The ORFs of the remaining genes are extended to cover all intergenic regions while maintaining the predicted translational frame.

#### Reference metamer calculation and compression

Metabuli computes reference metamers from the extended ORFs and their translations. Metabuli utilizes TANTAN (24) to mask low-complexity regions with repeat probability *>* 0.9 and excludes the masked regions from metamer calculation. All computed metamers are sorted numerically and then by their associated species ID. Because metamers encode amino acids in the leading significant bits, metamers encoding the same amino acid sequence are placed consecutively after sorting, and within them, they are grouped by codon usage, followed by their associated species ID. Then, redundant metamers from the same species are removed, retaining only one of them (Supp. Fig. 1c). The reduced metamer list is then further compressed as follows (Supp. Fig. 1d). The full numerical value of the first metamer is stored. For all other metamers on the list, only the increment value from the previous metamer is stored. The 64-bit encoding of the first metamer and the increments are then scanned as four slices of 15 bits each (the last four bits are unused). The slice of the least significant bits and any slice where some of the bits are turned on are copied and stored in 16 bits with one extra bit for an *end* flag. The end flag indicates whether the copied slice was the last one to be saved from a specific 64-bit value (where 1 = the copied slice is the last one). The optimal case is when only one slice is stored per metamer, yielding a compression ratio of four. The more reference metamers there are, the smaller the increments between consecutive ones tend to be, so the compression rate becomes closer to four. For example, when Metabuli was used to create a database from genomes of NCBI RefSeq release 217 (∼1.1TB), the compression rate was about three. Throughout this procedure, the reference sequence ID associated with each metamer is stored alongside it as well as information concerning metamer redundancy.

### Metabuli: Database decompression and usage

The values of the first metamer and the increments can be computed back from the stored compression by concatenating corresponding slices in a 64-bit data type. From the second metamer, their values are sequentially calculated by summing up each increment.

### Metabuli: Classification

#### Metamer match search

Query metamers from reads are sorted and compared to the reference metamer list to find matches (Supp. Fig. 1f-g). Because both query and reference metamers are sorted, a single iteration through the lists is enough to find all matches.

#### Calculating Hamming distance

After a query metamer is matched with reference metamers that are identical to it on the AA level, the closest matches are selected based on their DNA Hamming distance to the query. The distance between query and reference metamers is calculated using a Hamming distance lookup table (Supp. Fig. 2a-b). In this table, the 3-bit representations of any pair of synonymous codons are used as indices to retrieve their distance. The distances of a match are summed up when the total DNA Hamming distances of matches are compared to choose the closest metamer match (Supp. Fig. 2c).

#### Computing sequence similarity and assigning taxonomy

The matched metamers of each read are grouped by genus and species and examined by their coordinates on the read. For each species, only matches within a minimum of four consecutive matches are used to reduce the risk of random matches. Two matches are considered consecutive when 1) their query metamers are extracted from positions that differ by 3 nt in the same translational frame, and 2) the Hamming distances within the overlapping region are identical. Such matches to each genus are aligned to the query to compute the sequence similarity score between the query and the genus. The score is calculated based on the number of identical AAs, the Hamming distances, and the query length (Supp. Fig. 1h and Supp. Fig. 2c). Next, Metabuli assigns the read to the genus of the highest sequence similarity score. If more than one genus has scored the highest, the query is classified as the LCA of the best-scoring genera. Similarly, the matches found from the assigned genus are grouped by each species to assign the query to the species of the highest sequence similarity (Supp. Fig. 1i).

### Metabuli: Metabuli-P

Notably, as with other short k-mer-based classifiers, relying on few matches can often lead to false positive or overconfident classifications. False positive classification occurs mainly when the matched region is short. The similarity between a pair of sequences is expected to be higher if the pair belongs to the same lower taxonomic rank (rather than a higher rank). Over-confidence occurs when a read is classified at lower ranks like species or subspecies with not enough sequence similarity. To address this, Metabuli’s precision mode (Metabuli-P) uses two sequence similarity thresholds to avoid false and overconfident classifications. These thresholds were set based on similarity score distributions within prokaryotic and viral genera and species (Supp. Fig. 3).

#### Distribution of sequence similarity scores

We investigated the distribution of sequence similarities underlying TP and FP classifications using prokaryotes and viruses. Prokaryotic and viral species were identified based on two criteria: 1) there was at least one other species belonging to the same genus in the database, and 2) the database contained genomes of at least two of their subspecies. For prokaryotes, we could find 435 species, from the 22,819 GTDB genomes, that met the two criteria. We then designed two settings: subspecies-exclusion and species-exclusion. In the subspecies-exclusion, for each of the 435 species, one subspecies was included in the reference database while one of its sibling subspecies was excluded from it and used to simulate query reads. In the case of species-exclusion, the same database was used, and for each of the 435 species a random sibling species from the same genus was used to generate query reads. In both settings, 45,000 paired-end reads for each query genome were simulated using Mason2 as described above. In the case of viruses, we used NCBI taxonomy and Viral RefSeq. We could not find enough viral species fulfilling both criteria. Therefore, for the subspecies-exclusion setting, we applied the second criterion to find 211 species with at least two subspecies. In the case of the species-exclusion setting, the first criterion was applied to find 889 genera that have at least two species. In both settings, 10,000 paired-end reads were simulated from each query genome. Then, we used Metabuli to classify query reads in the various test settings and examined the sequence similarity scores underlying the TP or FP classifications.

#### Determining thresholds

Examination of the sequence similarity distributions revealed that FP’s relative frequency peaks under sequence similarity of 0.1 (Supp. Fig. 3). Furthermore, the vast majority: 89.2-99.7% of all TPs are associated with a sequence similarity score greater than 0.15 (Supp. Fig. 3a-d), while many FPs (29.4.3-59.7%) are associated with a lower score (Supp. Fig. 3e-h). Therefore, Metabuli-P is set to leave a query as unclassified if its best genus-level similarity score is lower than 0.15. In the subspecies-exclusion settings, 97.0% (prokaryote) and 82.2% (virus) of the TPs are associated with a similarity score greater than 0.5 while only 14.6% (prokaryote) and 57.4% (virus) of the FPs scored as high. Thus, Metabuli-P is set to classify a read at the species level or a lower rank only if it has a similarity score of *>* 0.5 to at least one species.

### Prokaryote benchmarks

#### Inclusion test

We examined the 22,819 complete genome or chromosome level assemblies in the GTDB by their species and identified 1,626 species that had at least two subspecies with a genome in the database. Of these, 435 species were used for the score threshold setting of Metabuli-P (Supp. Fig. 3). The remaining (1,191) contributed two subspecies each, from which 6,150 paired-end reads were simulated with Mason2 (∼15M reads in total). Each of the same genomes was also used to simulate ONT reads of 3X depth using PBSIM3. Performance metrics were measured at species and subspecies ranks. The genomes used for database creation and read simulation are listed in Supp. Tables.

#### Exclusion test

The 22,819 GTDB genomes were examined by their genera. We identified 802 genera, which had at least two species with a genome in the database. Of these, 435 were used for the score threshold setting of Metabuli-P (Supp. Fig. 3). The remaining 367 genera were used for the exclusion test. In this setting, ∼ 50,000 reads were simulated from each species (∼ 20M reads in total) using Mason2. PBSIM3 was used to simulate ONT reads of 3X depth from each of the species. Performance was measured at genus rank. The assembly accessions of the genomes for database creation and read simulation are available in Supp. Tables.

### Pathogen detection benchmarks

#### Inclusion test

Reference databases were built using genomes from NCBI Viral Refseq and five SARS-CoV-2 variant genomes (alpha, beta, delta, gamma, and omicron). We manually included these variants as children of SARS-CoV-2 to the NCBI taxonomy database. Two sets of RNA sequencing data from COVID-19 patients were used as query reads. One set was prepared from three patients infected by the beta variant (25), and the other from three patients infected by the omicron variant (26).

#### Exclusion test

The database for each tool was constructed using the taxonomy of NCBI and the genomes of Viral Ref-Seq, excluding all SARS-CoV-2 sequences. Due to this exclusion, SARS-CoV-1 is the closest relative in the reference database to any variant of SARS-CoV-2. RNA-seq data from thirteen SARS-CoV-2 patients and five controls prepared in a host-response study were used as query reads (27). The estimated number of SARS-CoV-2 reads in each sample was calculated by multiplying the total number of RNA-seq reads by the reads per million (RPM) of reads aligned to the SARS-CoV-2 genome. The RPM values were taken from the original study.

### CAMI 2 benchmarks

We used paired-end reads of strain-madness, marine, and plant-associated datasets and taxonomy provided in CAMI2 (15). In the case of CAMI2-provided reference databases for DNA- and AA-based tools (*nt* and *nr*), where there are no one-to-one relationships between their entries, it is possible to encounter under- or over-representation of some taxa. This discrepancy can lead to a potentially unfair comparison between the two groups of classifiers. To replace the CAMI2-provided databases, we used the reference genomes and proteomes in the prokaryote inclusion test. The references and a mapping from accessions to taxonomic IDs used in CAMI2 were provided to each classifier for database creation. Metabuli, Centrifuge, and KrakenUniq used 7,318 genomes, which together with two additional genomes were used for Kraken2, Kraken2X, Kaiju, and MMseqs2. CAMI2 provides 10, 21, and 100 query samples for the marine, plant-associated, and strain-madness benchmarks, respectively. To reduce the runtime of the benchmarks, we took all, every second, and every tenth query samples, respectively. We also used the CAMI2-provided ground truth labels for each read. When measuring performance at the species and genus ranks, we ignored classifications for reads whose ground truth taxon is at a higher rank than the rank of measurements.

### Benchmarks with real metagenomes

We challenged the classifiers on two distinct metagenomes: one obtained from a well-studied environment, specifically a human gut sample (SRR24315757), and the other from a less-studied environment, a marine sample (SRR23604821) (28). The same GTDB databases as in the prokaryote inclusion test were used.

### Resource measurement

Maximum RAM usage (maximum resident set size) and elapsed time of each tool were measured with the GNU `time -v` command. The average performance over five repeated measurements is reported (Supp. Table 1).

### Software versions and options

All benchmarks were performed with Kaiju v1.9, Kraken2 v2.1.2, KrakenUniq v0.7.3, Centrifuge v1.0.4, and MM-seqs2 v13.45111. We run Centrifuge with `-k 1` option to report at most one classification per read. For Kraken2, `--minimum-hit-groups` was set as 3 following a recommended usage (29). Struo v0.1.7 was used to download genomes and make taxonomy dump files for GTDB benchmarks. Mason_simulator v2.0.9 and PBSIM3 v3.0.0 were used to simulate query reads.

### Computing resource

For the resource measurement, we used a server and a MacBook Air. The server was equipped with a 64-core AMD EPYC 7742 CPU and 1TB of RAM, and the MacBook Air (2020) had 8GB RAM and an Apple M1 chip (8-core CPU with 4 performance cores and 4 efficiency cores). A server with 2 ×64-core AMD EPYC 7742 CPUs and 2TB of RAM was used for other benchmarks.

---

**Extended Data Fig. 1.**
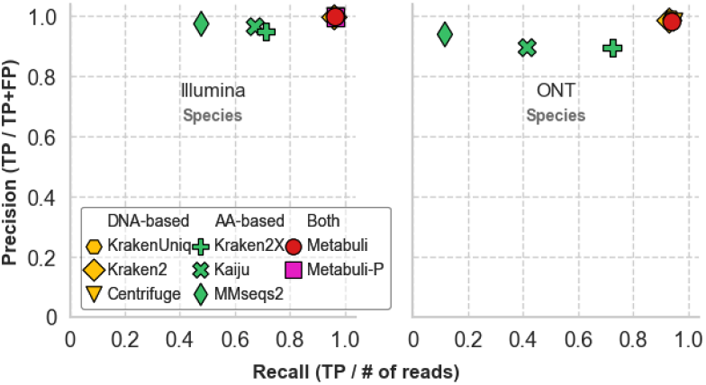
Prokaryote inclusion test results at species rank. Precision and recall of tools in the benchmarks of Fig. 2a were measured at species rank.

**Extended Data Fig. 2.**
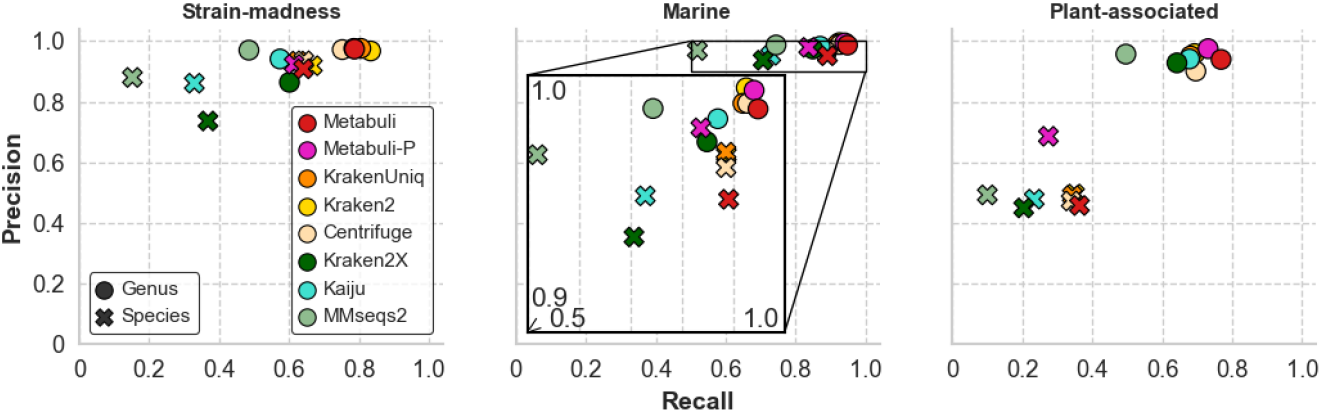
Benchmarks using CAMI2’s strain-madness and marine dataset. GTDB genomes and the CAMI2-provided taxonomy were used for reference construction. CAMI2-provided queries of strain-madness, marine, and plant-associated datasets were classified by each tool, and metrics were measured at the species and genus ranks.

**Extended Data Table 1.** Speed and memory usage

| Software | GTDB inclusion test |  |  | GTDB exclusion test |  |  |
| --- | --- | --- | --- | --- | --- | --- |
|  | DB size (GiB) | RAM (GiB) | Time (sec) | DB size (GiB) | RAM (GiB) | Time (sec) |
| Kraken2 | 43 | 44 | 24 | 41 | 42 | 55 |
| KrakenUniq | 309 | 306 | 169 | 294 | 292 | 272 |
| Centrifuge | 40 | 41 | 218 | 39 | 41 | 247 |
| Kraken2X | 11 | 12 | 26 | 10 | 12 | 47 |
| Kaiju | 39 | 41 | 145 | 38 | 41 | 582 |
| MMseqs2 | 37 | 174 | 6075 | 34 | 173 | 6606 |
| Metabuli 32GiB | 69 | 27 | 525 | 66 | 22 | 512 |
| Metabuli 64GiB | - | 47 | 484 | - | 37 | 465 |
| Metabuli 128GiB | - | 91 | 473 | - | 68 | 448 |
| Metabuli 256GiB | - | 173 | 480 | - | 129 | 450 |
| Metabuli MacBook 6GiB | - | 5 | 5640 | - | 4 | 6680 |

\* About 15M and 20M of 150nt paired-end reads were used in the inclusion and exclusion test
\* Metabuli has an `--max-ram` option that limits maximum RAM usage. Here, runs with the option set as 6, 32, 64, 128, or 256 GiB were presented.
\* All runs utilized 32 threads except for “Metabuli Macbook”, which used 8 threads.

**Supplementary Fig. 1.**
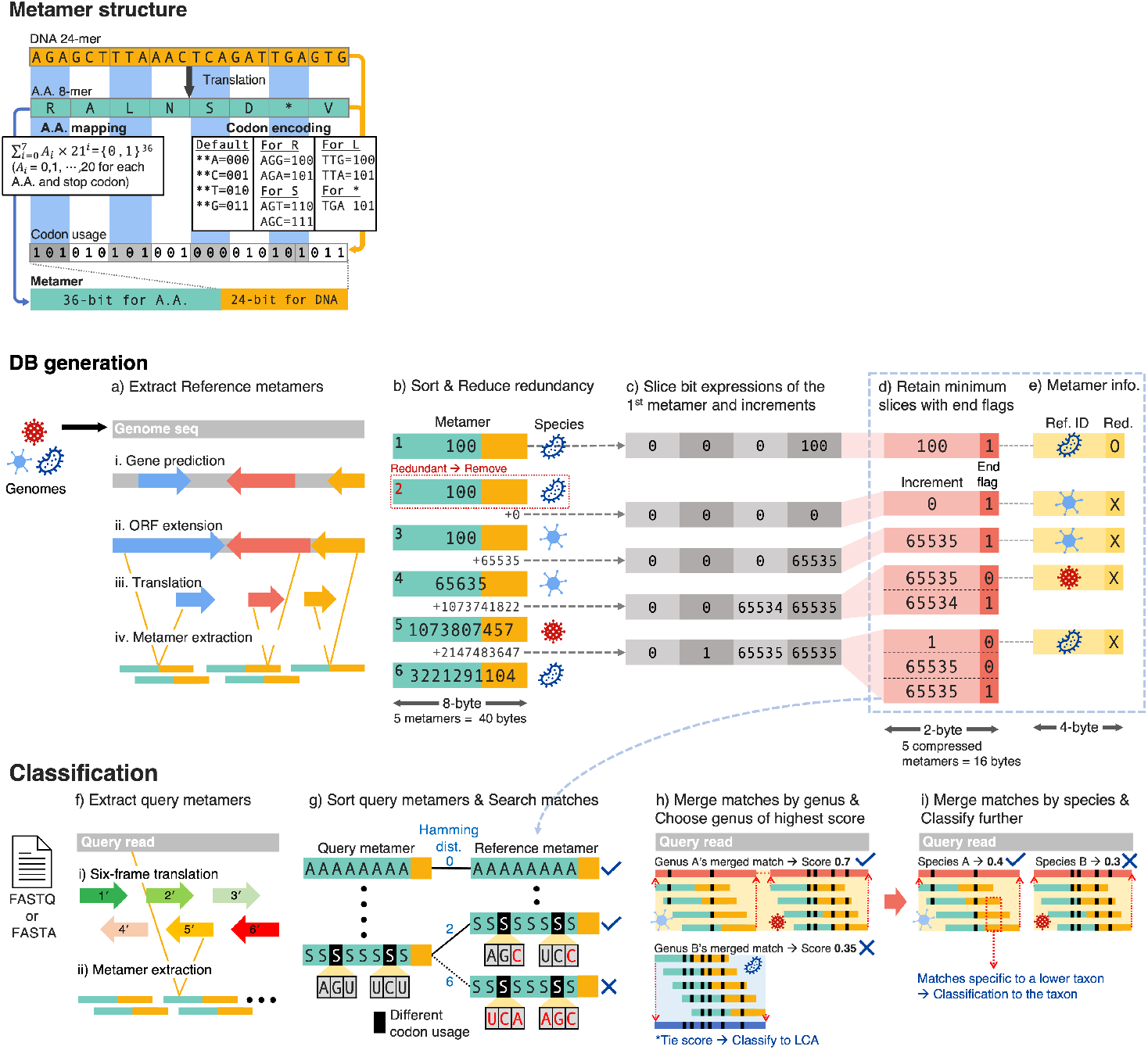
Metabuli’s workflow and metamer structure. Metamer structure. An AA 8-mer and the codon usage of each AA are stored in a metamer using 60 bits. The codon encoding table shows how synonymous codons of each AA are mapped to 3-bit encodings. **Database generation**. a) Metabuli builds a database from genomes in FASTA format. It predicts ORFs using Prodigal and extends them to cover intergenic regions. The extended ORFs and their translations are used to compute reference metamers. b) The computed metamers are sorted numerically and redundant ones from the same species are removed. c) The 60-bit expression of the first metamer and each difference (increment) between two consecutive metamers are scanned as four 15-bit slices. d) The slice of the least significant bits is stored, followed by all other non-zero slices. An end flag is added to each slice to indicate if it was the last one to be stored from the 60-bit expression. This allows grouping the slices by the 60-bit expression they correspond to. e) The reference ID and redundancy of each metamer are stored in a separate list. **Classification**. f) Metabuli takes query files in FASTA or FASTQ format. It scans each read in six frames and computes metamers from the DNA fragments and their translations. g) Query metamers are sorted and compared to reference metamers to find perfect AA matches. Among these, matches with the smallest DNA Hamming distance are selected (Supp. Fig. 2a-b). h-i) The matches of each genus are aligned to the query to score the genus, and then matches from the best genus are grouped by species to find the best species (Supp. Fig. 2c). Matches specific to a lower rank are used for lower-rank classifications.

**Supplementary Fig. 2.**
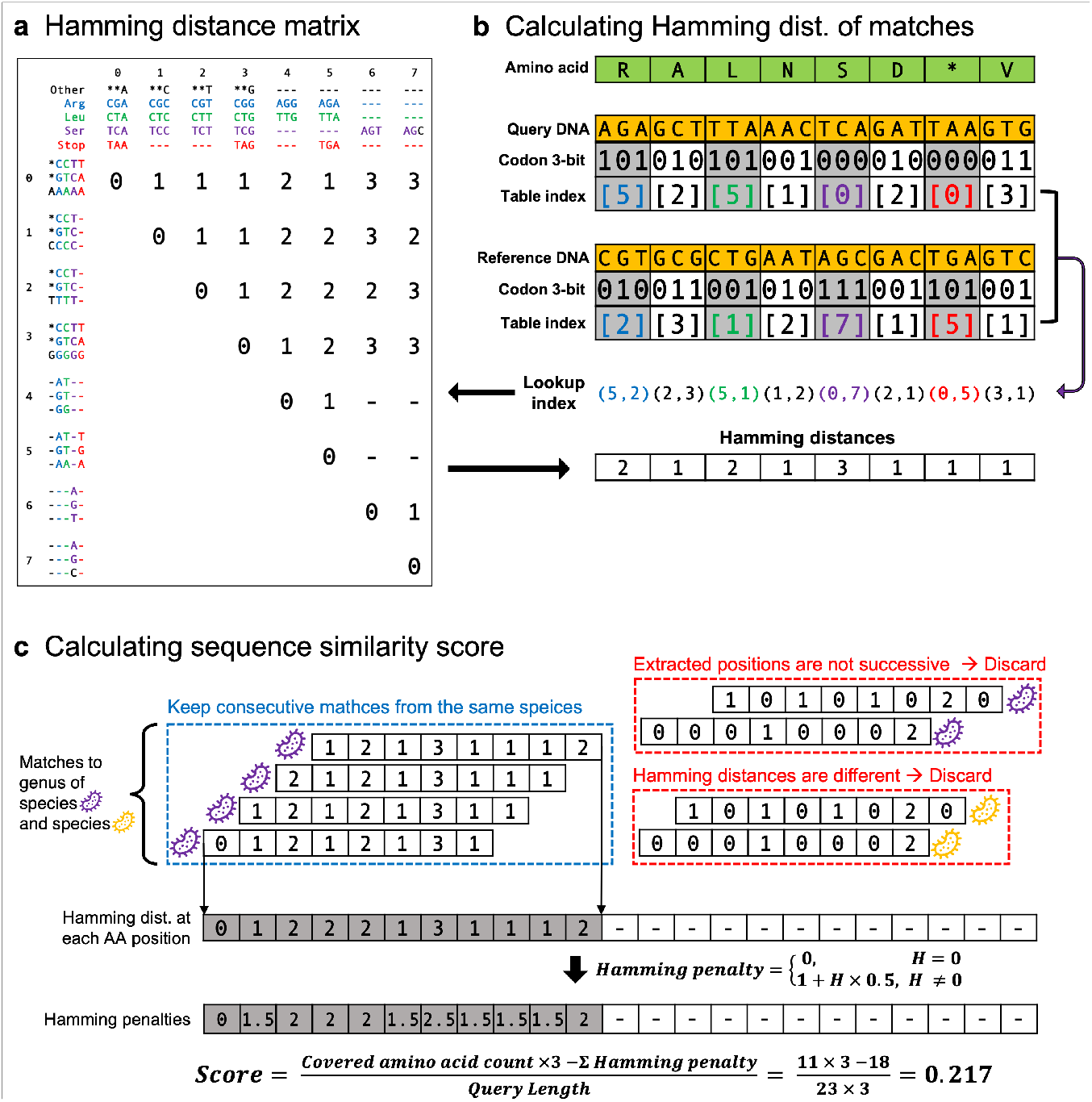
Calculating Hamming distance. **a)** The Hamming distance lookup table stores the distance between two codons of an identical AA pair in an 8 by 8 matrix. **b)** An example of Hamming distance calculation. An AA sequence (top, green) can be a translation of two different DNA sequences (DNA 1 and 2, orange). The 3-bit codon encodings for the same AA are used to index the Hamming distance lookup table. The Hamming distances of 8 codon pairs are summed up to get the total Hamming distance. **c)** Matches to a genus are aligned along the translated query, and Hamming distance at each position is pooled (See Supplementary Fig. 1g-h). The score of the taxon is calculated using the number of covered amino acids and Hamming penalty.

**Supplementary Fig. 3.**
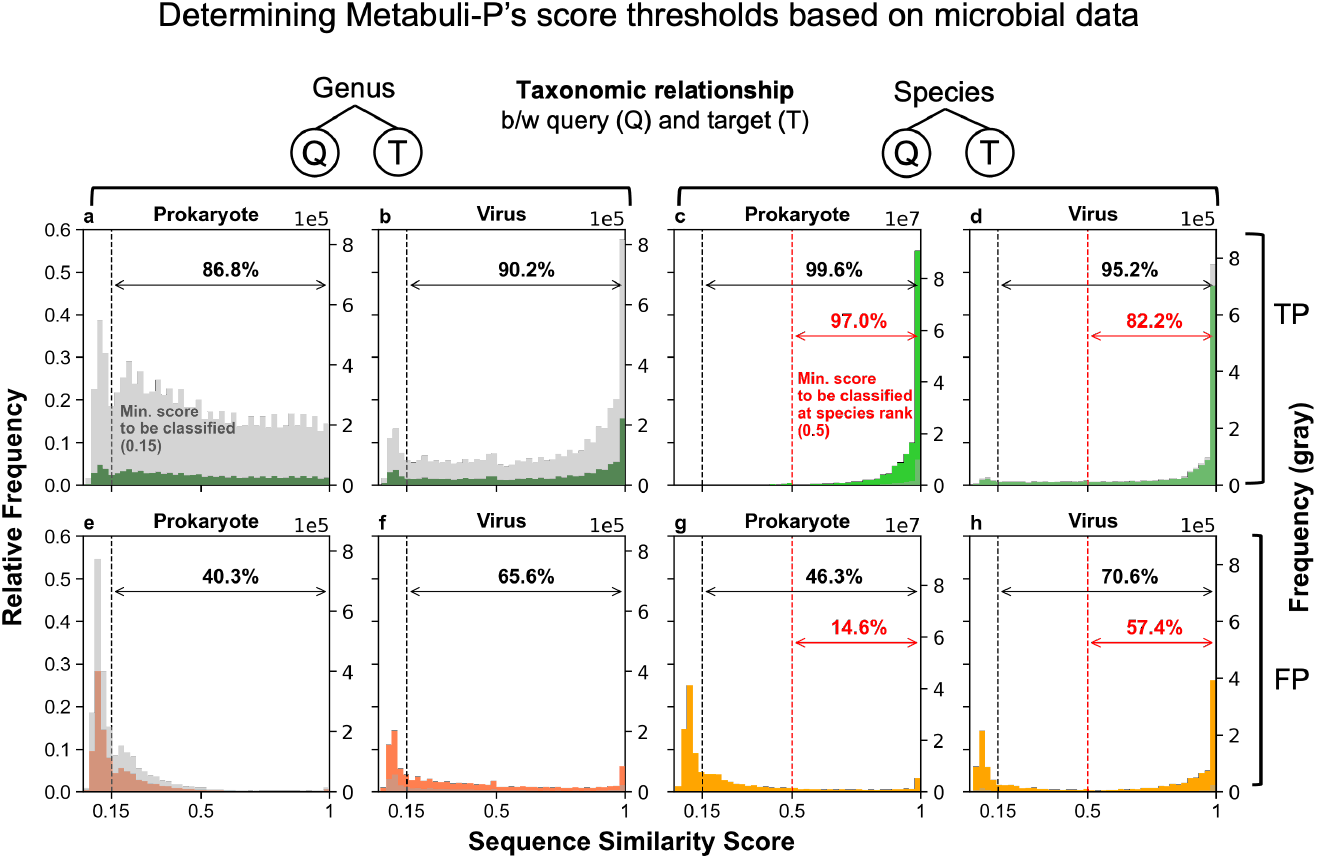
Sequence similarity score distribution. The distribution of sequence similarity scores was examined in prokaryotic and viral data (full details in Methods). The thresholds for classification are marked as dashed lines. These thresholds were selected because most TP classifications were made with sequence similarity that is greater than these thresholds, while many of the FPs have lower sequence similarity. **a-h)** Setting a threshold of 0.15 as the minimal sequence similarity for classification removes 53.7-59.7% of FP prokaryotic classifications and 29.4-34.4% of the viral FPs while retaining 86.8-99.6% of all TPs. **c-d)** Out of species-level classifications, 97.0% (prokaryote) and 82.2% (virus) of TPs have sequence similarity score > 0.5. So Metabuli-P has a threshold of 0.5 as the minimal sequence similarity for species-level classification to avoid over-confident low-rank classification. A similar threshold could not be identified for the genus level (a-b).

## Notes

https://github.com/steineggerlab/Metabuli

### Competing Interest Statement

The authors have declared no competing interest.

### Summary of Updates

Benchmarks using simulated long reads were added. Masking low-complexity regions during database creation. Code and data availability sections.

